# Liver physiological T1rho dynamics associated with age and gender

**DOI:** 10.1101/172478

**Authors:** Yì Xiáng J Wáng, Min Deng, Jiang Lin, Anthony WL Kwok, Eric KW Liu, Weitian Chen

**Author notes:** Correspondence to: Dr. Yì Xiáng Wáng. Department of Imaging and Interventional Radiology, Faculty of Medicine, The Chinese University of Hong Kong, New Territories, Hong Kong SAR.

## Abstract

**Purpose:** Using a single breathhold black blood sequence, the current study aims to understand the physiological ranges of liver T1rho relaxation for women and men.

**Materials and Methods:** This volunteer study was conducted with institutional ethics committee approval, and included 62 females (age mean: 38.9 years; range: 18-75 years) and 34 males (age mean: 44.7 years, range: 24-80 years). MRI was conducted with a 3.0 T scanner, with six spin-lock times of 0, 10, 20, 25, 35, 50msec and a single breathhold of 12 seconds. Six slices were acquired for each examination.

**Results:** Female liver T1rho value ranged between 35.07 to 51.97ms, showed an age-dependent decrease with younger women had a higher measurement. Male Liver T1rho values ranged between 34.94 to 43.39 ms, and there was no evidential age-dependence. For females, there was a trend that liver T1rho value could be 4%-5% lower during menstrual phase than nonmenstrual phase. For both females and males, no evidential association was seen between body mass index and liver T1rho.

**Conclusion:** Liver T1rho physiological value for males have relatively narrow distribution, while physiological value for females have wider distribution, and decreases with age.

**Key points:** 1. Liver T1rho shows an age-dependency in women, with young women showing higher measurement. This age-dependency of liver T1rho measurement is not evidential in men. Post-menopausal women have similar liver T1rho value as men.

2. Women at menstrual phase may have slight lower liver T1rho measurement.

3. No association was noted between body mass index and liver T1rho

4. When blood signal suppression sequence is used, in a population of 62 healthy women and 34 healthy men, the highest measured liver T1rho was 52 msec for young women, 44.7 msec for post-menopausal women, and 43.4 msec for men.

---

Chronic liver disease is a major public health problem worldwide [1,2,3]. The epidemic trend of chronic liver disease is expected to increase owing to an aging population, the growing epidemic of obesity and non-alcoholic steatohepatitis (NASH). Liver biopsy remains the standard of reference for the diagnosis and staging of liver fibrosis. However, it is an invasive procedure with potential complications. Histologic assessment of fibrosis is also an inherently subjective process and subject to sampling variability. Both animal experiments and clinical studies have revealed that liver fibrosis, even early cirrhosis, is reversible. Earlier stage liver fibrosis is more amenable to therapeutic intervention. Treatment with combined therapies on underlying etiology and fibrosis simultaneously might expedite the regression of liver fibrosis and promote liver regeneration [4, 5]. Even when the underlying etiology of liver fibrosis could not be eradicated, therapies on liver fibrosis might help restrict the disease progression to cirrhosis. A noninvasive, objective and quantitative technique for assessing liver fibrosis and monitoring disease progression or therapeutic intervention is highly desirable.

The spin-lock radiofrequency (RF) pulse used in T1rho imaging introduces the sensitivity to chemical exchange effect, which makes it feasible to use T1rho to probe macromolecular environment of human tissue [6,7]. Wang *et al* and Zhao *et al* described the usefulness of T1rho MR imaging for assessment in rat liver fibrosis models caused by biliary duct ligation and carbon tetrachloride intoxication, suggesting increased T1rho value is associated with collagen deposition in the fibrotic liver but less so with inflammatory edema [8,9]. Studies in human subjects, both in healthy volunteers and liver fibrosis patients, have since been reported [10-19]. With a two-dimensional fast gradient echo sequence, Deng *et al* and Zhao *et al* reported the mean T1rho value was 42.8±2.1 ms (B_0_=3T, B_1_=500Hz) in healthy volunteers [10-12]. Allkemper *et al* [14] reported that normal liver T1rho value was 40.9±6 2.9 ms (B_0_=1.5T, B_1_=500Hz), and elevated liver T1rho measurement was significantly associated with the Child-Pugh staging of the patients. In patients with chronic liver diseases, Takayama *et al* [16] demonstrated liver T1rho value showed significant positive correlations with the serum levels of total bilirubin, direct bilirubin, and indocyanine green (ICG-R15), and significant negative correlations with the serum levels of albumin and γ-glutamyl transpeptidase. Xie *et al* [19] reported fatty liver did not affect the diagnostic performance of T1rho for detecting liver fibrosis, and suggested a upper limited of 49.5ms for non-fibrotic livers.

To overcome the artifacts associated with respiration motion during T1rho image acquisition [20], Singh *et al* proposed a pulse sequence enabling acquisition of single slice 2D T1rho MR imaging with four different TSLs in a single breathhold of 12 seconds without suppression of blood signal [18], or 16 seconds with single inversion recovery for fluid suppression [21]. More recently Chen *et al* proposed a 2D black blood T1rho MR Relaxometry technique for liver imaging [22]. Black blood effect of 2D fast spin echo (FSE) sequence and double inversion recovery were utilized to achieve blood signal suppression. In a pilot volunteer study, Wáng *et al* [23] demonstrated that compared with multiple breathhold bright blood signal technique, this new sequence reduces image artifacts, improve image quality, and improves scan-rescan reproducibility, with promise for clinical application. Using this single breathhold black blood sequence, the current study aims to understand the physiological ranges of liver T1rho relaxation for women and men. This knowledge is critically important for developing liver T1rho as a clinical tool for practical application.

## Materials and Methods

This volunteer study was conducted with the approval of the institutional ethics committee, and informed consents were obtained. One hundred healthy volunteers without known history of liver diseases were recruited from the local community via advertisement. Two obese participants, with one subject had a BMI of 33.8 and one subject had a weight of 99 kg, failed the examination. T1rho value measurement could not be precisely performed in one participant due to multiple cysts in liver. MRI showed one subject had liver solid tumor, but follow-up was lost and therefore excluded in final analysis. During the course of this study decision was made to recruit more females than males, as females showed greater age-dependence. Finally, there were 96 volunteer participants, including 62 females (age mean: 38.9 years; range: 18-75 years) and 34 males (age mean: 44.7 years, range: 24-80 years).

All participants underwent T1 weighted and T2 weighted liver anatomical imaging, and single-breathhold black blood FSE liver T1rho imaging [22, 23], using a Philips Achieva TX 3.0 T scanner equipped with dual transmitter (Philips Healthcare, Best, the Netherlands). The subjects were scanned in supine position. A 32 channel cardiac coil (Invivo Corp, Gainesville, USA) was used as the receiver and body coil was used as the transmitter.

2D axial images were acquired with phase encoding along anterior-posterior direction. Radiofrequency (RF) shimming was applied to reduce B1 inhomogeneity (Fig 1). A RF pulse cluster that can achieve simultaneous compensation of B1 RF and B0 field inhomogeneity was used for T1rho preparation [24, 25]. The parameters for MR imaging included: TR/TE 2,500/15 msec, in-plane resolution 1.5 mm × 1.5 mm, slice thickness 6 mm, SENSE acceleration factor 2, half scan factor (partial Fourier) 0.6, number of signal averaging 1, delay time for SPAIR 250 msec, delay time for double inversion recovery 720 msec, spin-lock frequency 500Hz. Images with six spin-lock times (TSLs) 0, 10, 20, 25, 35, 50msec were acquired, with a single breathhold of 12 seconds. Six slices were acquired for each examination. Breathhold was trained for the volunteers before the scan started. A good breathhold is essential to obtain high quality T1rho map (see supplementary document 1).

**Fig 1.**
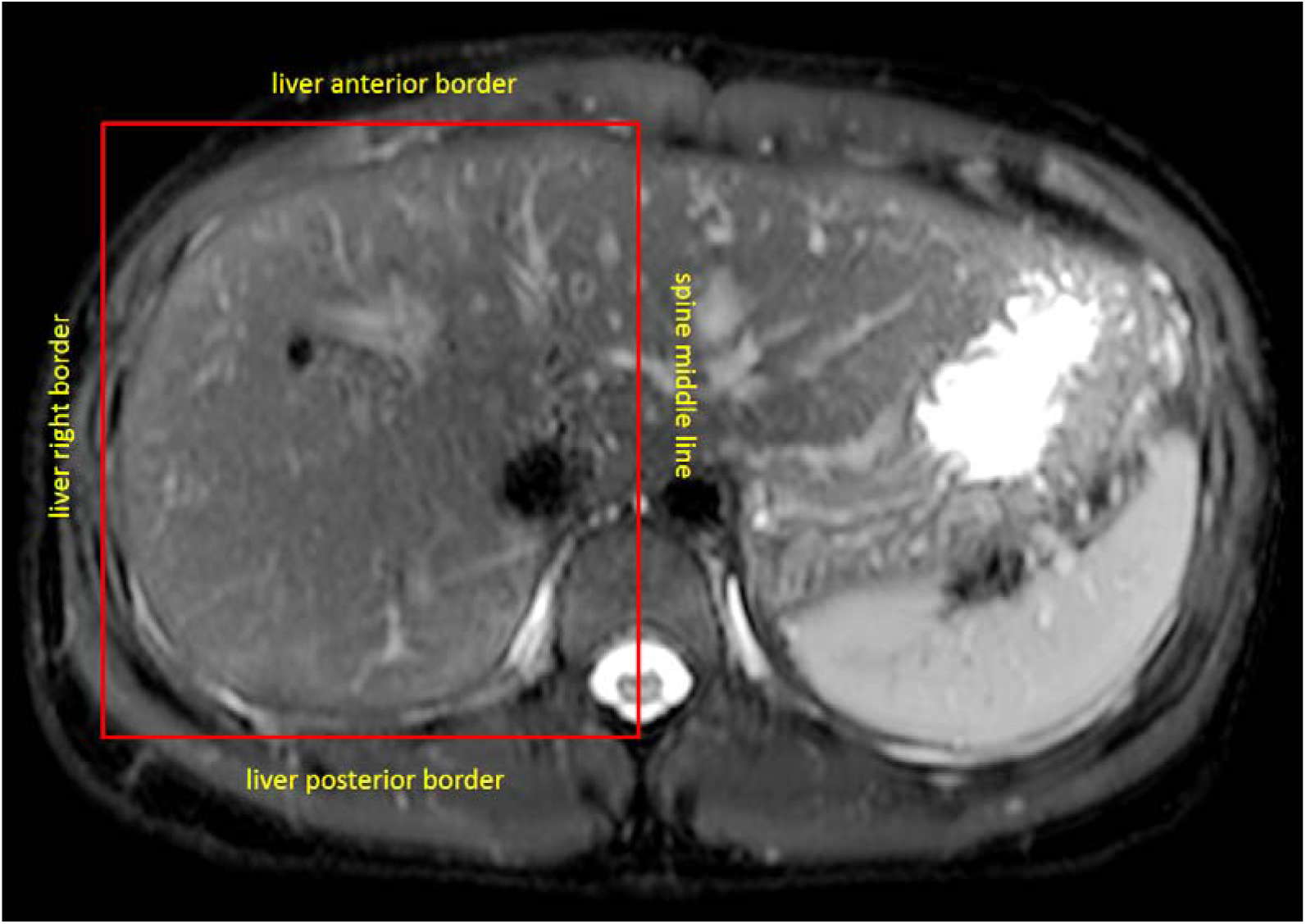
A shimming box placed on T2 weighted image. This shimming box is defined by spine middle line, anterior, posterior, and right liver borders.

All images were processed using Matlab R2015a (Mathworks, USA). T1rho maps were computed by using a mono-exponential decay model, as described by the following equation: *M* (TSL) = *A* · exp (- TSL/T1 rho), where *A* is a constant scaling factor and TSL is the time of spin-lock. Non-linear least square fit with the Levenberg-Marquardt algorithm was applied. Maps of coefficient of determination (R2) were also generated for the evaluation of goodness of fit. Only T1rho values for pixels associated with R^2^>0.80 were included in the subsequent region of interest (ROI) placement and T1rho analysis to eliminate the unreliable poorly fitted T1rho values due to artefacts [10, 11]. With the aim to standardize measurement, a histogram protocol was used, and the pixel in highest three bars were included as the liver T1rho reading (Fig 2). It was shown that histogram approach and ROI approach give very similar measurement [12].

**Fig 2.**
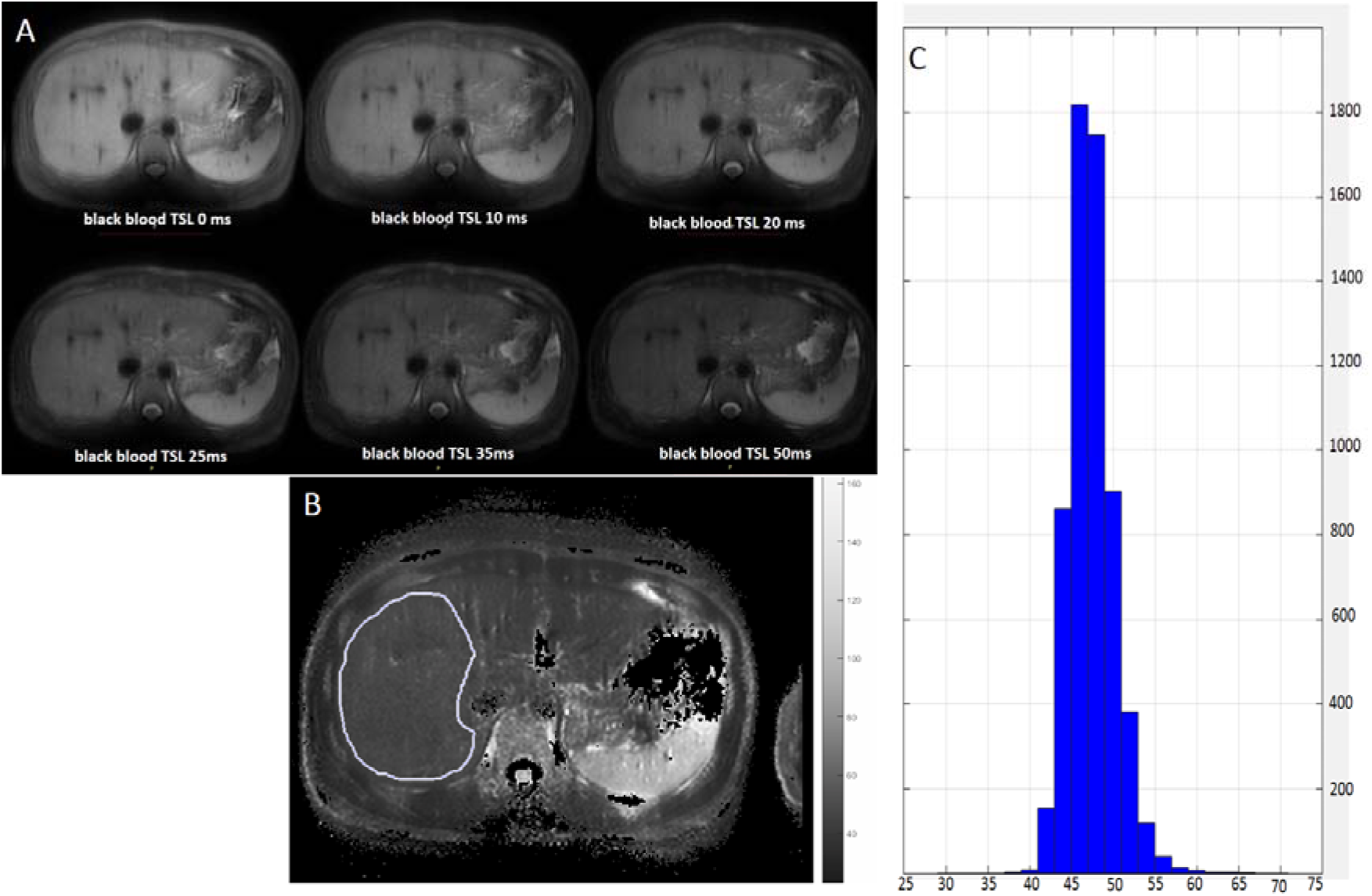
A: T1rho images with TSL (spin lock time) of 0, 10, 20, 25, 35, 50 ms. B: T1rho map and a ROI drawn on liver parenchyma. C: Histogram analysis of the ROI drawn in B.

During the course of this study, decision was made to include body mass index (BMI) information, and menstrual cycle information for females. The body mass index (BMI) information was obtained in 51 female participants and 24 male participants. The menstrual cycle information was obtained in 37 female participants. Five females were scanned twice, at menstrual phase and second half of the non-menstrual phase, to investigate the menstrual cycle’s impact. Our pilot study showed the impact of meals on liver T1rho values was minimal, and therefore this point was not considered (see supplementary document 2 for more information). Liver T1rho values in males and females were separately analyzed.

An abdominal radiologist had more than 15 years of experience read all the standard anatomical T1 weighted and T2 weighted fat suppressed images. Incidental findings in the liver and biliary system included the following: liver cysts in 30 participants, liver hemangioma (or cysts) in 2 participants, gallbladder adenomyosis in 3 participants, biliary stone in 4 participants, cholecystitis (thickened gallbladder wall) in 1 participant, biliary hamartoma 1 participant, and small lymph node seen in porta hilar in 1 participant. These incidental findings, as well as other incidental findings in pancreas, spleen, and kidneys were communicated to the participants. Further follow-ups or other examinations such as ultrasound were recommended when appropriate.

## Results

Physiological liver T1rho values are shown in Table 1 and Fig 3, 4. The box-and-whisker-plot presentation is shown in Fig 5. Female Liver T1rho values showed with a strong age-dependent decrease (Fig 3, s*lope* of estimated linear fit =-0.17, p<0.0001). There was no evidential age-dependence of male liver T1rho values (s*lope* of estimated linear fit = -0.17, p>0.05). Post-menopausal females had similar liver T1rho distribution as males. The lowest liver T1rho measurements were very similar for females and males (table 1, Fig 5).

**Table 1.**
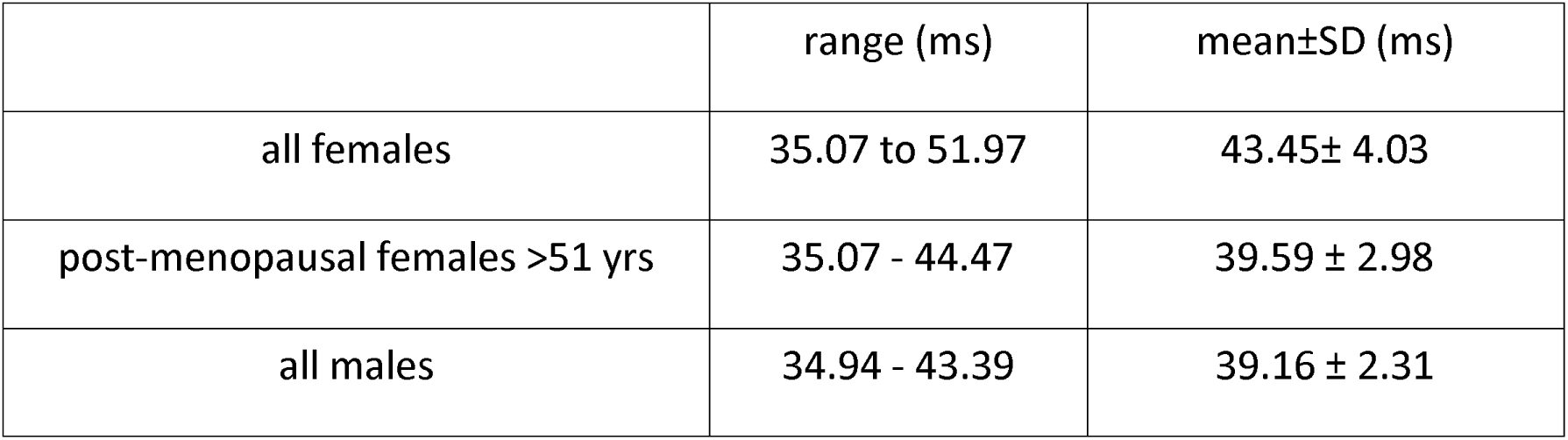
Physiological liver T1rho values of female and male subjects

**Fig 3.**
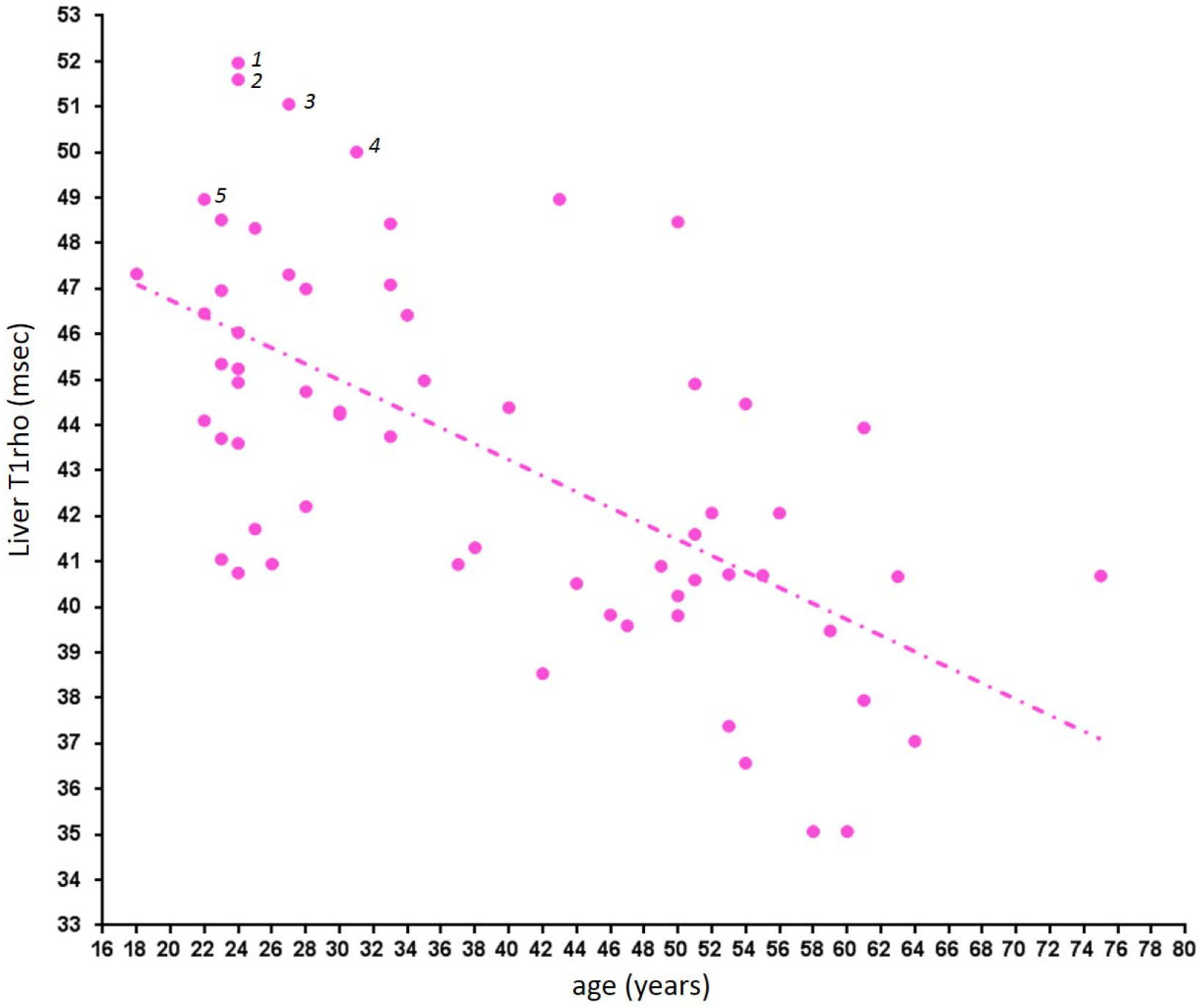
Liver T1rho in female subjects. An age-dependent decrease is noted. Cases 1 & 3: blood liver function and viral hepatitis-b antigen tested negative; cases 2 & 4: blood liver function, viral hepatitis-b antigen, and viral hepatitis-c antibody tested negative; case 5: blood viral hepatitis-b antigen tested negative.

**Fig 4.**
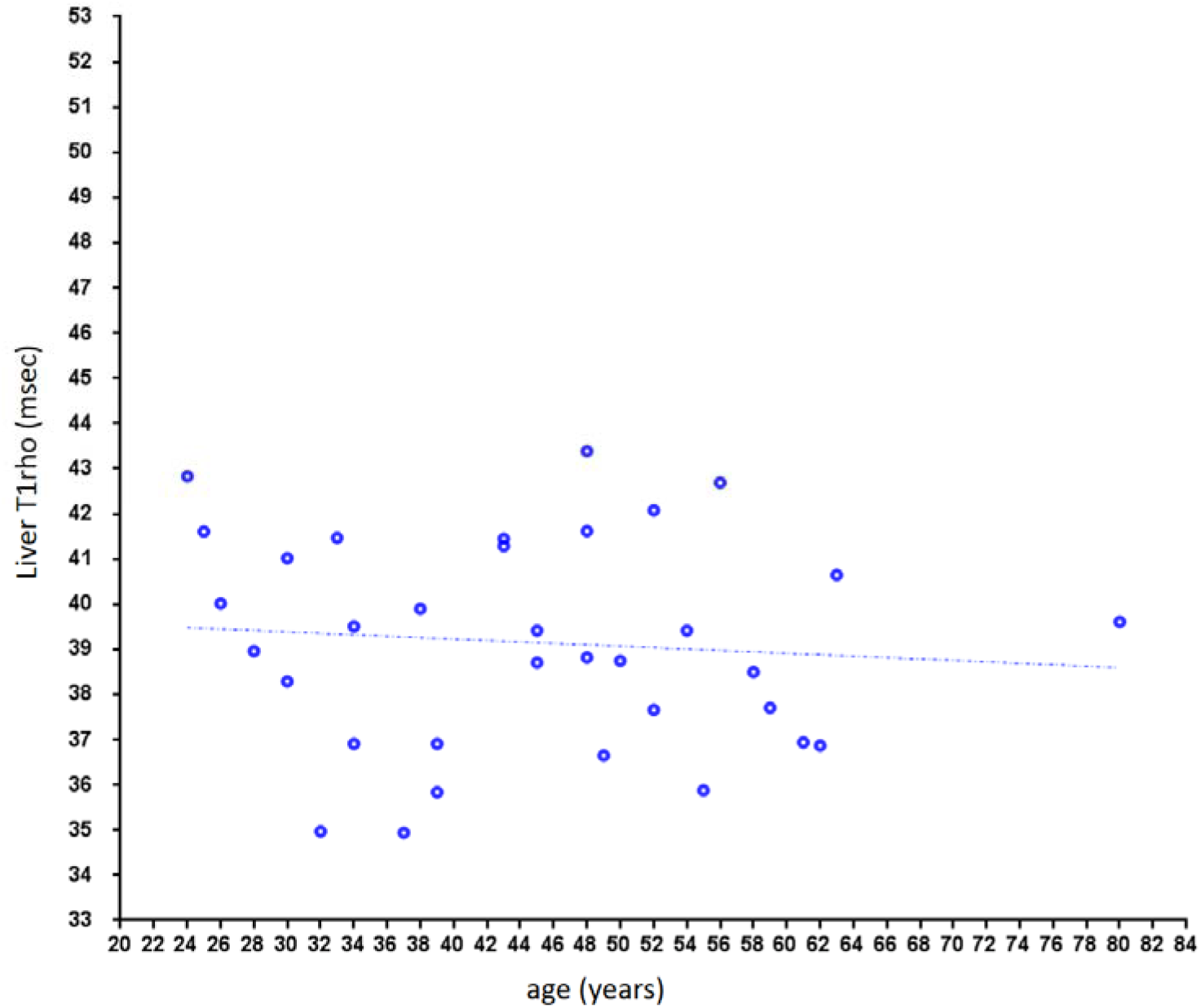
Liver T1rho in male subjects. No notable age-dependency is observed.

**Fig 5.**
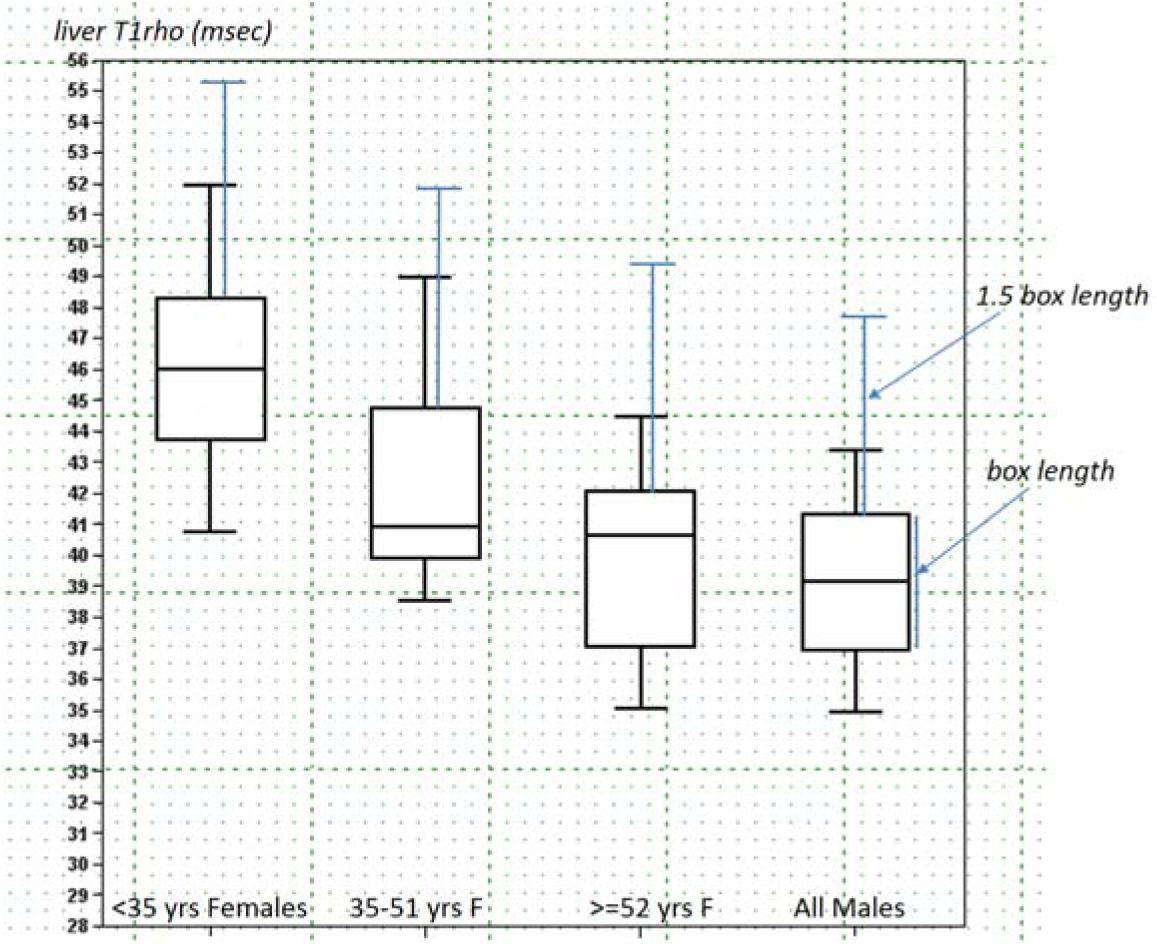
Box-whisker plots of liver T1rho of three female age groups of <35 yrs, 35-51 yrs, and ≥ yrs, and males of all ages. The 1.5 box length defined upper limits are denoted in blue lines (supplementary document 3).

The menstrual cycle information of 37 female participants were recorded in this study. Thirty six participants were in non-menstrual phase, with mean liver T1rho value being 45.69±3.64 ms. Eleven participants were in menstrual phase, with liver T1rho value being 43.53±3.49 ms (4.7% lower than menstrual phase subjects, *p*= 0.053 for one-sided T-test). For the five subject scanned at menstrual phase and second half of the non-menstrual phase, liver T1rho was 42.74±4.31 ms and 44.52 ±2.36 ms respectively, with menstrual phase being 4.2% lower (*p*=0.07 for one-sided T-test, Wilcoxon signed ranks test *p*=0.14, Wilcoxon signed ranks test).

The relationship between liver T1rho and BMI is shown in Fig 6. For both females and males, no evidential association was seen between liver T1rho and BMI.

**Fig 6.**
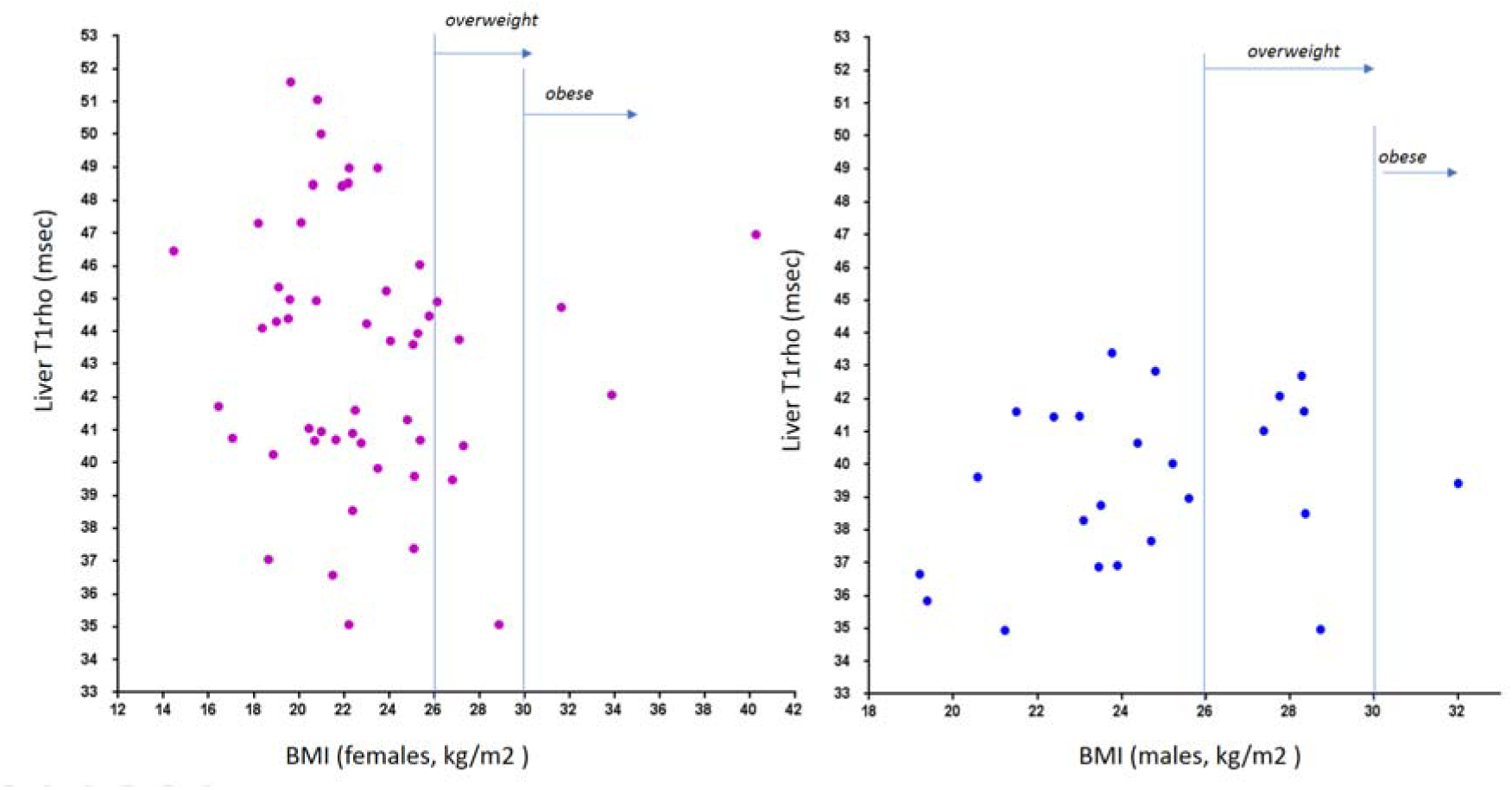
Relationship between liver T1rho and BMI. No notable association is observed.

## Discussion

T1rho has the potential to be a sensitive marker of liver diseases. This has been indicated in the earlier animal studies [8, 9, 26]. In a recently study, Koon *et al* [27] reported on day-10 after biliary duct ligation all experimental animals can be separated from normal liver based on T1rho measurement, when liver histology resembled early stage fibrosis, and fibrous area counted for 0.38%±0.44% of liver parenchyma area. Liver T1rho may also be used as marker of liver function, as demonstrated by Allkemper *et al* [14] and Takayama *et al* [16]. With a gradient echo bright blood based sequence, recently Xie *et al* suggested the upper limited of physiological T1rho measurement may be 49.5 ms in humans [19], which is higher than the results of Allkemper *et al* [14]. More recently, with a limited cohort of 20 subjects Wáng *et al*’s reported that physiological range of liver T1rho may be age- and gender- specific [23]. However, till now the physiological range of liver T1rho has not been systematically studied.

The advantage of the single-breathhold black blood fast spin echo based sequence has been reported [23]. Single breath-hold reduces T1rho imaging’s susceptibility to motion displacement occurring at different spin lock times. Fast spin echo based black-blood T1rho measurement is also less contaminated by blood signal [23]. The current study successfully implemented a ‘protocol-driven’ study of liver in 96 volunteer subjects. To avoid occasional operational failures, 6 slices were acquired for each subject and liver T1rho was measured with a histogram approach so to avoid potential operator induced bias. The breath-hold duration was 12 seconds per slice acquisition, and was well tolerated by all participants.

The liver T1rho normal range will be relatively easier to establish for male subjects as they likely distribute in a narrow range (Fig 4). In the study of Koon *et al*. [26] with 32 young male rats (200-250 gram), the physiological liver T1rho value was 38.38±1.53 ms (range: 36.05~41.53 msec), which is very similar to the human data obtained in this study (mean±SD: 39.16 ± 2.31 ms, range: 34.94 ms to 43.39 ms). The range is slightly wider in human results is likely to due to laboratory animals were more homogeneous in their physiological condition. Since male liver T1rho value is more consistent across ages, it is more appropriate to compare liver T1rho by black blood fast spin echo sequence and bright blood gradient echo sequence using male subject data, with the mean difference being 3.2 ms (39.16 ± 2.31 ms vs 42.36±2.00 ms) [10, 11]. The physiological range of liver T1rho for female subjects demonstrated strong age-dependence, with younger females having higher T1rho measurement. Actually, to our surprise, liver T1rho can be higher than 50 ms in younger females, actually such high measurement was considered to be associated with liver fibrosis in previous studies [14, 19]. Indeed, we asked all subjects with liver T1rho > 50 ms to have liver function test and at least viral hepatitis-b test, which turned out to be negative for all of them. On the other hand, post-menopausal females older than 51 years had T1rho values similar to male subjects. According to Box-Whisker analysis, if data with 1.5 box length from the upper quantile are considered as outliers, then the upper limit for females age groups of <35 yrs, 35-51 yrs, and ≥51 yrs would be 55.20 ms, 52.05 ms, and 49.60 ms respectively, and the upper limit for males would be 47.93 ms. However, we expect the real upper limits would be smaller than these values. This needs to be confirmed in future studies.

The fact that age-dependent liver T1rho is strong in females, but not apparent in male, led us to investigate the possible liver T1rho value changes during menstrual cycle, as we expect liver T1rho is influenced by female sex hormones. For the five subjects scanned both at menstrual phase and second half of the non-menstrual phase, liver T1rho during menstrual phase was 4.2% lower. In addition, 11 participants scanned in menstrual phase had a mean liver T1rho value 4.7% lower than other 36 participants scanned in non-menstrual phase. Our reading to these results is that females at menstrual phase may have slightly lower liver T1rho, which may be in the order of 4%-5% lower than non-menstrual phase. While statistical power can be increased with access to larger sample sizes, we expect this may not be very relevant for most subjects’ clinical care. For marginal cases, liver T1rho imaging at non-menstrual phase may improve consistency. Our preliminary analysis showed liver T1rho has no evidential association with BMI. The World Health Organization defines overweight as a BMI of 25 or higher and obesity as a BMI of 30 or higher. For Chinese adults aged 20 years or older, recently it was reported that a BMI of 24.0–25.9 for men and women was deemed as optimal [28, 29]. Xie *et al* recently reported that fatty liver itself does not affect liver T1rho quantification [19]. Allkemper *et al*’s data also showed T1rho did not correlate significantly with degree of steatosis of liver [14] The ability to demonstrate a male-female difference, as well as the age-dependence in females, demonstrates the sensitivity of T1rho analysis. The liver plays a pivotal role in metabolism and xenobiotic detoxification. Instead of being static, liver is an organ full of physiological dynamics, for example, liver mass in birds and humans show diurnal fluctuations [30-32]. Recently Sinturel *et al* showed that that, in mice, liver mass, hepatocyte size, and protein levels follow a daily rhythm, whose amplitude depends on both feeding-fasting and light-dark cycles [33]. Estrogen exhibits a number of important roles in the body, such as to promote coagulation, aid in maintaining proper fluid balance, and foster increases in high density lipoproteins and decreases in low density lipoproteins that lead to favorable lipid profiles. Estrogen has been shown to regulate the structure and function of mitochondria, particularly in tissues that have a high energy demand. Mitochondria have an important role in ATP production, cell metabolism and apoptosis, and homeostasis of reactive oxygen species as well as hepatocyte metabolism of glucose, lipids, and proteins [34]. Hepatic estrogen receptors mediate estrogen action in the liver, and estradiol has a favorable role in chronic liver disease, which is suggestive of a protective effect of estrogens against NAFLD (Non-alcoholic fatty liver disease) in women [35]. There are observations of persistent increases in the incidence of NAFLD in women beyond middle age, whereas such continued increases in NAFLD incidence are not demonstrated in men [36, 37]. Aging increases the risk for NAFLD in premenopausal women [37, 38]. Age is a risk factor for NAFLD in women, but not in men [38]. The physiological gender difference in blood system has been also noted such as men and women have different mean haemoglobin levels in healthy venous blood, with the mean level of women approximately 12% lower than that of men [39, 40]. The gender- and age-related dependences observed with liver T1rho may be related to till now unknown biophysics of liver tissue characteristics which are at least partially regulated estrogen.

Another point to be noted is that T1rho value is dependent on the sequence design [41], therefore comparison of T1rho values measured by different acquisition methods may not be directly compared. While the T1rho imaging method described in this study is practical for clinical application, we also expect conversion factors can be developed for comparison of T1rho sequences with different preparation designs.

There are a number of limitations for this study. The study participant number is small, as we did not include patients with mild liver diseases in the study, the upper limit of normal liver T1rho remain undecided. The data obtained in this study can provide background information for future patient studies. For example, a man with liver T1rho > 45 ms should raise high suspicion; on the other hand, a young women with liver T1rho = 50 ms may be normal. BMI and menstrual cycle information were not obtained in all participants, as the decision to include these information was made during the course of the study, rather than pre-planned. However, the inclusion of BMI and menstrual cycle information in this study can be considered as ‘random sampling’. Due to practical difficulty, the study into menstrual cycle’s impact on liver T1rho remains preliminary. Ideally, liver T1rho imaging should be performed for each phases of menstrual cycle. However, the results from this study showed the magnitude of difference between different phases would be small. Another possibility will be that not all the ‘healthy’ volunteers might be actually healthy. However, all the subjects with liver T1rho > 50 ms had liver function and viral hepatitis-b checked to be negative; and the liver T1rho for males has relative narrow distribution and also very similar to data from previous experimental studies. A full understanding of the physio-pathological meaning of liver T1rho is expected to require multiple studies involving more healthy volunteers as well as patients of different etiologies. Another technical limitation of current T1rho sequence is insufficient RF pulse energy for T1rho imaging for some obese patients. Further sequence optimization is required to overcome this difficulty.

In conclusion, this study reports the normal liver T1rho value in healthy volunteers. The physiological value for males has relative narrow distribution, while physiological value for females has wider distribution, and decreases with age. In females the liver T1rho measurement tends to be 4-5% lower during menstruation phase. Post-menopausal women have similar liver T1rho range as men.

## Acknowledgement

This study was partially supported by grants from the Research Grants Council of Hong Kong SAR (Project nos. 476313 and SEG CUHK02).

## Supplementary documents

### Supplementary document-1: Breathhold training

A good breathhold is important for obtaining motion-free liver T1ro data. Breathhold is trained for the scan subjects before the scan starts. It is more likely that the diaphragm and liver position will shift if we ask the scan subjects to hold their breath after full end-inspiration or full end-expiration. Therefore, scan subjects are asked to hold their breath during usual-depth breathing. After hearing the ‘hold-breath’ instruction, the subjects are given time to allow diaphragm back to the most comfortable position. A time delay was allowed between the scan operator to give ‘hold-breath’ instruction and to push the MR data acquisition start button, so that the volunteers had time to react to the ‘hold-breath’ instruction.

The respiration-gating balloon is placed on the top of the scan subjects’ upper abdomen, and the quality of the ‘breathhold’ is monitored on the respiration-triggering screen on the MRI console.

For MRI units with busy schedule, it is adviceable that the breathhold training is conducted before the scan subjects enter the MRI room. In this case, a mock training set of a simple combination of a balloon and a monitor will be useful.

### Supplementary document-2: Liver T1rho fasted and postprandial difference

There are a number of physiological changes in liver after a subject eats a meal, including increased blood volume and glycogen level. Therefore there is possibility that there will be difference of T1rho in fasted liver and post-prandial liver.

The previous study with multi-breath gradient echo sequence did not demonstrate statistically significant difference between fasted group and post-meal group [Br J Radiol. 2012;85(1017):e590-5]. Liver T1rho value was 43.1±1.4ms (n=9 subjects) in the fast status and 43.0±2.4 ms (n=13 subjects) in the post meal status (p=0.867). The results of individual subjects who were scanned twice are shown as below (table S1).

**Table S1.** Gradient echo multiple S-breath fast echo sequence measured fasted and postprandial liver T1rho relaxation. Note the scans were not performed in the same day.

|  | T1 rho_fasted (ms) | T1 rho_postprandial (ms) | difference |
| --- | --- | --- | --- |
| case 1 | 45.2 | 44.9 | -0.66 % |
| case 2 | 47.3 | 43.4 | -8.25% |
| case 3 | 41.8 | 39.3 | -5.98% |
| case 4 | 41.0 | 38.6 | -5.85% |
| case 5 | 44.8 | 42.3 | -5.58% |
| case 6 | 42.4 | 41.6 | -1.89% |
| case 7 | 42.3 | 39.8 | -5.91% |
| case 8 | 42 | 42.7 | +1.67% |
| case 10 | 44.2 | 44.3 | +0.23% |
| *mean* | *43.44* | *41.88* | *-3.58%* |

Since we expect the single-breath black blood sequence can increase the sensitivity of T1rho imaging, we tested three subjects in the same day fasted group and post-meal, and the results are as below (table S2):

**Table S2.**
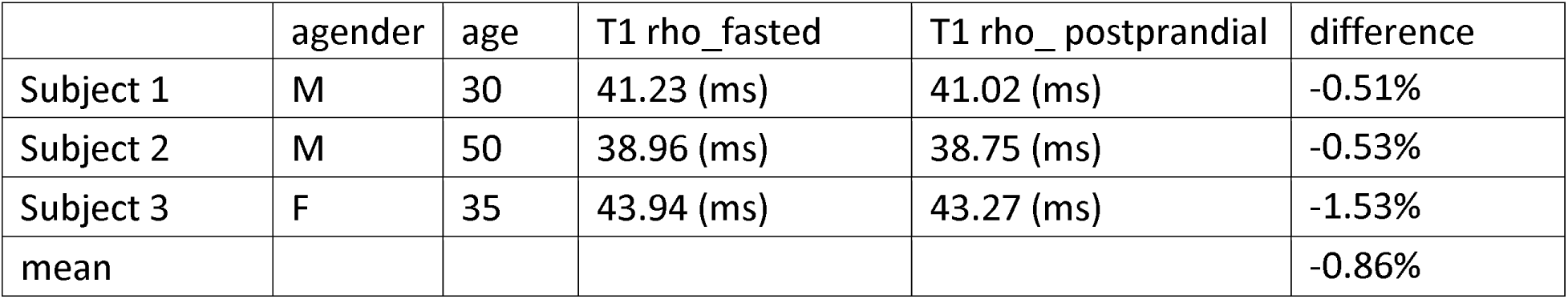
Single-breath fast echo sequence measured fasted and postprandial liver T1rho relaxation. Paired measurement was performed in the same day.

|  | agender | age | T1 rho_fasted | T1 rho_ postprandial | difference |
| --- | --- | --- | --- | --- | --- |
| Subject 1 | M | 30 | 41.23 (ms) | 41.02 (ms) | -0.51% |
| Subject 2 | M | 50 | 38.96 (ms) | 38.75 (ms) | -0.53% |
| Subject 3 | F | 35 | 43.94 (ms) | 43.27 (ms) | -1.53% |
| mean |  |  |  |  | -0.86% |

Based on the these results, we conclude that post-meal liver has lower measurement of T1rho than fasted liver, but the difference was very small, and would not have clinical implications with single-breath black blood sequence.

### Supplementary document-3: Box-Whisker analysis of liver T1rho value

The Box-Whisker analysis of liver T1rho value is shown in Table S3 and Fig S1 as below. If data with 1.5 box length from the upper quantile are considered as outlier, the statistically the upper limit for females age groups of <35 yrs, 35-51 yrs, and ≥51 yrs is 55.20 ms, 52.05 ms, and 49.60 ms respectively, and the upper limit for males is 47.93 ms. Though we expect the real upper limit would be smaller than these values.

|  |  | median | Q1 | Q3 | box_length | 1.5 box_length | lower limit | upper limit |
| --- | --- | --- | --- | --- | --- | --- | --- | --- |
| females | <35 yrs | 46.04 | 43.75 | 48.33 | 4.58 | 6.87 | 36.88 | 55.20 |
|  | 35-51 yrs | 40.92 | 39.94 | 44.78 | 4.85 | 7.27 | 32.67 | 52.05 |
| | $\geq 52$ yrs | 40.67 | 37.05 | 42.07 | 5.02 | 7.53 | 29.52 | 49.60 |
| males | all ages | 39.19 | 36.93 | 41.33 | 4.40 | 6.60 | 30.34 | 47.93 |

Q1: lower quartile; Q2: upper quartile

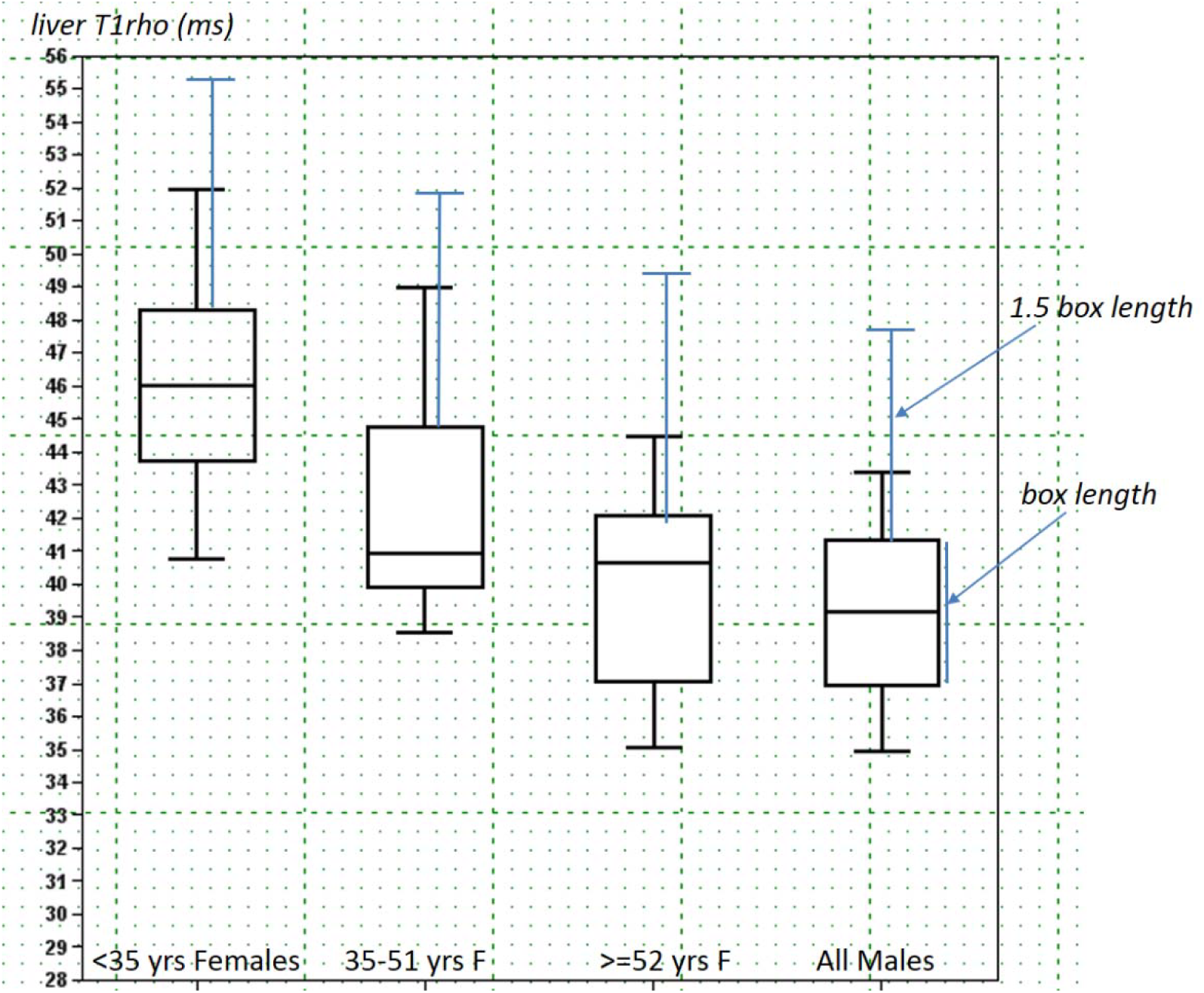

